# Modeling CNS inflammatory demyelination using myelin-directed CAR T cells

**DOI:** 10.64898/2026.11.16.767139

**Authors:** Christian Cordano, Sasha Gupta, Caroline Guglielmetti, Chanelle Shepherd, Madison Heady, Jason Duecker, Ahmed Abdelhak, Ben Ng, Robert Zhu, Wei Yu, Sharon A. Sagan, David G. Stefanoff, Jung H. Sin, Alexandra Beaudry-Richard, Angelica Montevirgen, Liana N. Kozanno, Stephen L. Hauser, Michael R. Wilson, Scott S. Zamvil, Ari J. Green, Wendell A. Lim, Milos S. Simic

## Abstract

Multiple sclerosis therapies remain confined to immunomodulatory agents that reduce inflammatory activity; none restores lost myelin, and disability can continue to accumulate independently of relapses. Development of remyelinating therapies has been hindered by models that incompletely reproduce the persistently inflamed lesions in which remyelination fails. Here we show that chimeric antigen receptor (CAR) T cells targeting myelin oligodendrocyte glycoprotein (MOG) build such lesions on demand. Human CD4+ CAR T cells transferred into immunodeficient mice infiltrated the brain, spinal cord, and optic nerve, producing perivascular demyelinating lesions, with reproducible optic nerve involvement. Within these discrete optic nerve lesions, treatment with the remyelinating compound clemastine reduced lesion size and improved optic nerve conduction, despite persistent inflammation. Thus, this MOG CAR T model reproduces key features of inflammatory demyelinating disease and enables testing of remyelinating therapies; more broadly, it introduces CAR T cells as programmable perturbative agents for building defined pathology in vivo.

## INTRODUCTION

Multiple sclerosis (MS) is a central nervous system (CNS) autoimmune disease characterized by focal demyelination caused by host immune cells and subsequent neurodegeneration^1–3^. When the demyelinating lesion acutely occurs along the optic nerve, this leads to a condition known as optic neuritis (ON). While there have been significant advances in immunomodulatory therapies to prevent new MS relapses, remyelination of existing lesions is an unrealized therapeutic goal in the treatment of MS. Addressing this gap is crucial, as chronic demyelination of axons is believed to lead to secondary axonal injury and contribute to irreversible neuronal loss and potentially progressive disability in MS^4^.

To further advance the field of remyelination and therapeutic prevention of neuroaxonal degeneration, improved animal models are imperative. Therapeutic development can significantly benefit from reliable and reproducible preclinical animal models that recapitulate critical features of human pathophysiology. The lack of such models is an obstacle in translating preclinical observations into clinical applications^5,6^. Decades of research have led to the development of diverse experimental autoimmune encephalomyelitis (EAE) murine models, the current mainstay animal model for demyelinating disease, ranging from inducible immunization models to spontaneous genetic models. Still, none of these mouse models present discrete demyelinating lesions within the CNS microenvironment, as is observed in human disease^7,8^. This critical deficit in pathological fidelity of EAE prevents its use to address the crucial question of whether remyelinating compounds can promote the formation of new myelin within discrete lesions^9,10^.

Here, we have developed a novel murine model for neuroinflammation-induced demyelinating disease using chimeric antigen receptor (CAR) T cells targeting the myelin sheath. CARs are synthetic receptors consisting of an extracellular domain that recognizes a surface-expressed antigen in an MHC-independent manner and an intracellular domain containing a signaling domain from the T cell receptor (CD3ζ) and a costimulatory signaling domain (traditionally CD28 or 4-1BB)^11,12^. T cells expressing the CAR directly attack cells expressing the target extracellular antigen. Anti-B-cell CAR T cells have revolutionized the treatment of hematopoietic malignancies and, more recently, are being explored for autoimmune conditions^12^. However, given CAR T cells’ potency, specificity, and engineerability, we hypothesized that we could leverage CAR technology to engineer cells that create a targeted demyelinating disease in animals. Rather than attempting to model the initiating immunological mechanisms of neuroinflammatory disease, CARs targeted towards a myelin-specific antigen could create a model of disease that results in a more faithful replication of the pathology of human disease by recapitulating the characteristic myelin injury seen in human disorders such as MS. This approach would provide a platform better suited for evaluating remyelinating therapies.

To target myelin and induce demyelination, we engineered human CD4+ T cells to express a CAR specific for myelin oligodendrocyte glycoprotein (MOG), a surface autoantigen of oligodendrocytes and myelin. First, we validated that the engineered T cells could cause a potent inflammatory response in vitro. We then injected these primary human T cells into immunocompromised mice. These T cells trafficked to the CNS and reproducibly caused encephalitis with focal clinical manifestations and optic neuritis with pathology showing prominent discrete demyelinating lesions. The use of immunocompromised mice eliminated the variability associated with host immune responses, enabling us to focus on the demyelinating effects of the CAR T cells. We then showed that clemastine, a drug with well-described remyelinating potential^13^, can induce remyelination of these discrete lesions. Thus, this study introduces a new class of mouse models that faithfully recapitulate focal demyelinating pathology and provide a robust platform for testing remyelinating therapies. More broadly, it demonstrates the potential of using engineered immune cells as programmable and modular tools for generating precise and reproducible disease models beyond their current use for therapeutic applications.

## RESULTS

### CD4+ T cells can be engineered to elicit an inflammatory response against a myelin-specific antigen in vitro

Fig. 1a illustrates our rationale to develop myelin-targeting T cells: these cells would traffic to the CNS and induce an inflammatory response when engaging with the myelin, creating a new model of demyelination. To that effect, primary human CD4+ T cells were engineered to express a CAR targeting MOG (myelin oligodendrocyte glycoprotein), a component of the myelin sheath known as a target in demyelinating diseases. We first sought to validate α-MOG CAR CD4+ T cell function in vitro. We established a test target cell line by engineering K562 cells to express full-length murine MOG (these cells are referred to as K562 MOG^+^). α-MOG CAR CD4+ T cells were co-cultured with the parental K562 or K562 MOG^+^ cells. Using flow cytometry, we confirmed that α-MOG CAR CD4+ T cells were specifically activated in the presence of K562 MOG^+^ through the increased expression of CD25, an activation marker, after 24h of co-culture (Fig. 1b i). These α-MOG CAR CD4+ T cells also showed significant proliferation, detected by the dilution of CellTrace^TM^ membrane dye over three days of co-culture with K562 MOG^+^ cells. CellTrace^TM^ is a membrane dye that allows for tracking cell proliferation: as the initially stained cell divides, the membrane dye is diluted in the daughter cells over time.

**Fig. 1:**
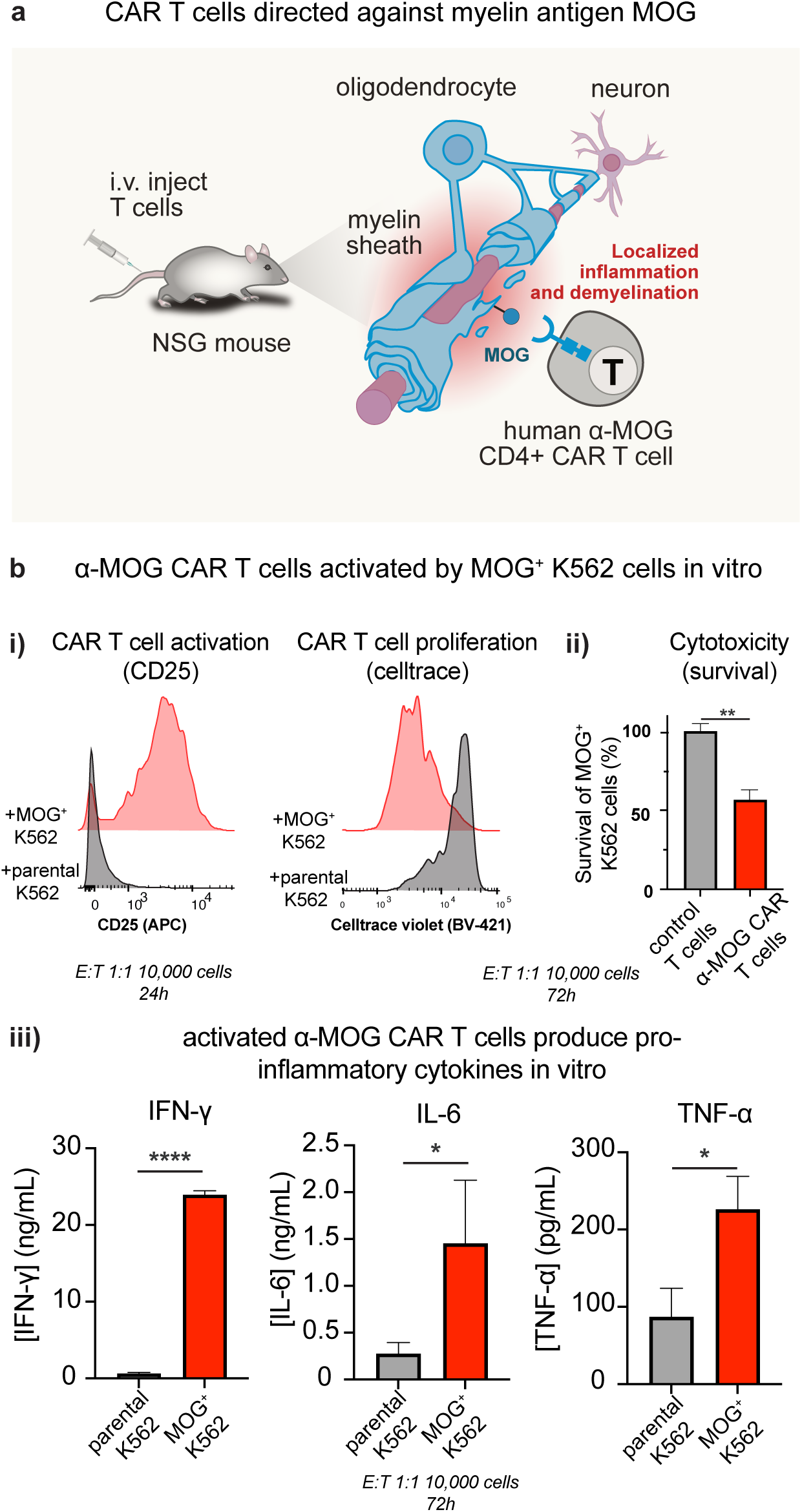
α-MOG CAR CD4+ T cells recognize MOG-expressing cells in vitro, resulting in T cell activation, proliferation, and secretion of inflammatory cytokines. a, Conceptual rationale for the design of directed immune responses to the myelin sheath using human T cells to model CNS demyelinating inflammation in a mouse. Human CD4+ T cells are engineered with a CAR that can recognize a CNS-specific myelin surface antigen, such as myelin oligodendrocyte glycoprotein (MOG). These cells are injected into immunocompromised NSG mice and trigger inflammation and demyelination in MOG-expressing tissues. b, Activation, proliferation, cytotoxicity, and pro-inflammatory cytokine secretion in vitro. Primary human CD4^+^ T cells transduced with α-MOG CAR were co-cultured with K562 priming cells, either expressing or not expressing MOG, at indicated E:T ratio, durations, and cell numbers. Activation state was assessed by flow cytometry using a CD25 stain. Proliferation was assessed by CellTrace^TM^ violet dilution assay. Cytotoxicity was assessed by MOG^+^ K562 survival. Representative histograms are shown. Secretion of pro-inflammatory cytokines IFN-γ, IL-6 and TNF-α was assessed by ELISA from the supernatant. n = 3, mean +/- std. Unpaired t-test.

To assess the cytotoxic potential of the α-MOG CAR CD4+ T cells, we evaluated the survival of the K562 MOG^+^ cells when co-cultured with either α-MOG CAR CD4+ T cells or control (untransduced) T cells. Only α-MOG CAR CD4+ T cells induced significant cytotoxicity over 72h of co-culture (Fig. 1b ii). To evaluate the inflammatory potential of the α-MOG CAR CD4+ T cells, we collected the supernatants and assessed them by ELISA for pro-inflammatory cytokines. We found that significant levels of IFN-γ, IL-6, and TNF-α were induced specifically in the presence of K562 MOG^+^ cells (Fig. 1b iii). The high levels of IFN-γ led us to assess if α-MOG CAR CD4+ T cells were predominantly exhibiting a Th1 response. We used intracellular staining against IFN-γ (specific for the Th1 subset) and IL-17A (specific for the Th17 subset) and found that α-MOG CAR CD4+ T cells had increased levels of intracellular IFN-γ but not IL-17A, consistent with a Th1-like response (Extended Data Fig. 1). When cultured with target K562 MOG^+^ cells, α-MOG CAR CD4+ T cells adopted a terminally differentiated phenotype with an increased percentage of effector memory RA (re-expressing CD45RA) (CD45RA^+^, CD62L^-^) cells, and upregulated surface-expressed exhaustion markers TIM3, PD1, and LAG3 (Extended Data Fig. 2).

Overall, α-MOG CAR CD4+ T cells showed a potent Th1 skewed inflammatory response specific to the presence of K562 MOG^+^ cells in vitro.

### Myelin targeting CD4+ T cells elicit a potent CNS-localized inflammatory response

Equipped with MOG-specific CAR T cells, we tested whether these cells could induce a localized inflammatory response in a mouse. We injected 5 x 10^6^ human primary α-MOG CAR CD4+ T cells into immunocompromised NSG (NOD.Cg-*Prkdc^scid^ Il2rg^tm1Wjl^*/SzJ) mice. The CAR T cells were also engineered to co-express green fluorescent protein (GFP) and firefly luciferase (effLuc) to allow for localization studies. We used α-human CD19 CAR CD4+ T cells as an antigen-irrelevant control, since this CAR does not recognize mouse CD19 or other mouse epitopes, and therefore cannot be engaged by any antigen in the recipient mice (Fig. 2a). Mice injected with α-MOG CAR CD4+ T cells suffered from various neurologic sequelae, ranging from paraparesis to paraplegia at various locations (forelimb, hindlimb, torso, or less often limp tail), and ataxia. These symptoms did not progress in an ascending stereotyped fashion as observed in EAE but occurred in a variable manner. Inevitably disease accumulated and led to progression in severity to meet euthanasia criteria, typically 2-3 weeks after injection. Weight loss paralleled mobility loss (Extended Data Fig. 3). Using longitudinal bioluminescence imaging, we observed an increasing signal in the heads of mice injected with α-MOG CAR CD4+ T cells compared to α-CD19 CAR CD4+ T cells, consistent with preferential localization and proliferation of the α-MOG CAR CD4+ T cells in MOG-rich areas (Fig. 2b). On MRI, mice showed increased ventricular volumes as shown by T2 weighted MRI, suggesting neuronal loss^14^ (Fig. 2c). Consistent with this observation, the serum levels of neurofilament light chain (NfL), a marker for neuronal damage^15–17^, were increased in α-MOG CAR CD4+ T cell-injected mice (p < 0.01) (Fig. 2d). CNS isolation of the mononuclear infiltrates showed high numbers of α-MOG CAR CD4+ T cells. Flow analysis of the isolated α-MOG CAR CD4+ T cells from CNS and spleen showed specific activation in the CNS compared to the spleen, as shown by CD25 staining (Fig. 2e). Further phenotyping indicated CNS-specific polarization of the α-MOG CAR CD4+ T cells towards a Th1 phenotype (Fig. 2f) consistent with the in vitro data (Fig. 1b iii and Extended Data Fig. 1).

**Fig. 2:**
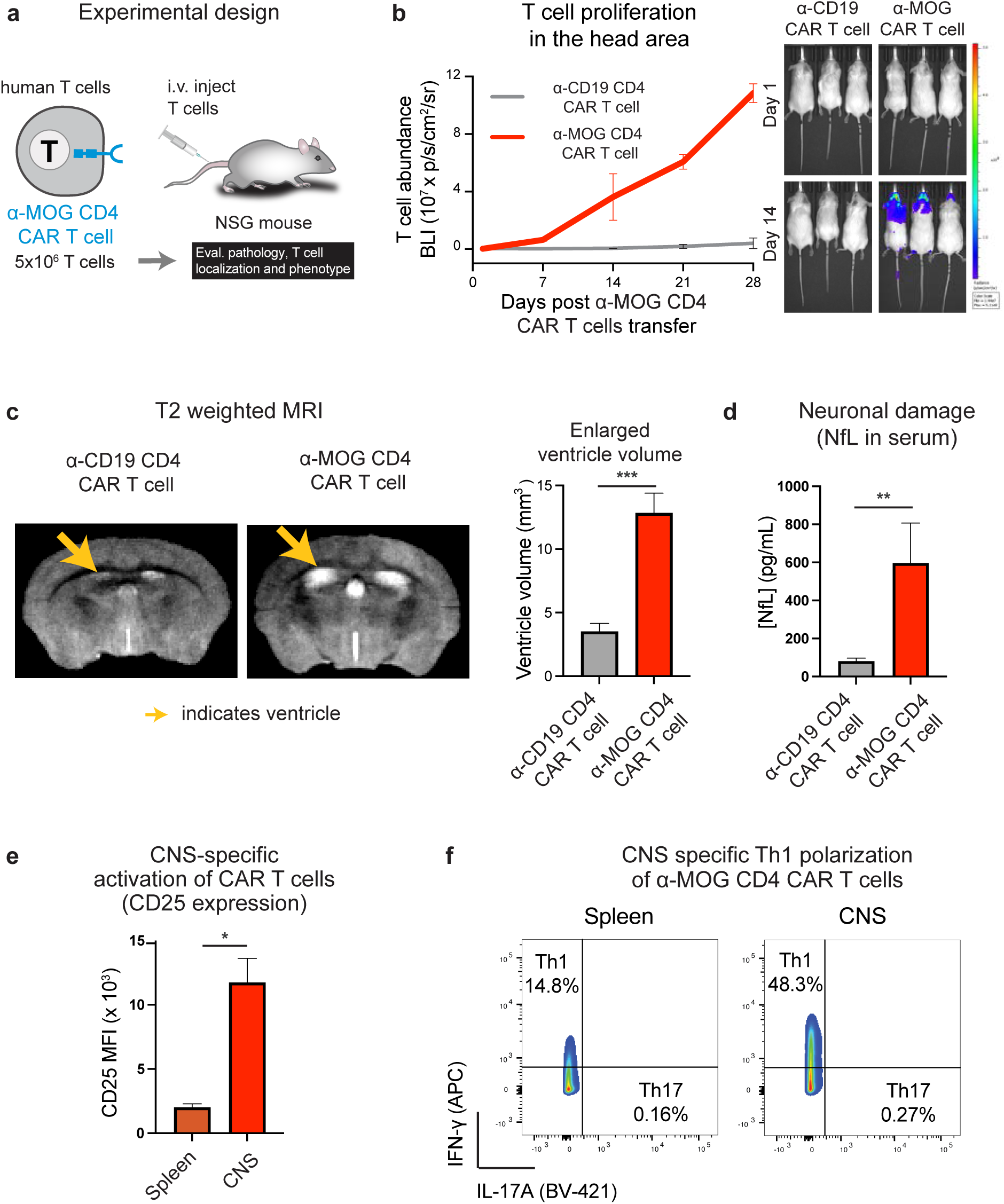
Human α-MOG CAR CD4+ T cells injected into mice localize to the central nervous system, proliferate and induce encephalitis with evidence of neuronal injury. **a**, In vivo experimental design of myelin-targeted CAR T cells. Primary human CD4+ T cells were engineered to express α-MOG CAR or control α-CD19 CAR, which does not cross-react with murine CD19. To allow tracking of the T cells both by fluorophore and bioluminescence, we co-transduced them to express GFP-2A-effLuc. 5 x 10^6^ primary human CD4+ T cells were injected intravenously (i.v.) in immunocompromised NSG mice and followed for evaluation (n=3). **b**, In vivo expansion of α-MOG CAR T cells. T cell location and expansion were followed over time by bioluminescence imaging. The bioluminescence from the head area was quantified. Representative images of Day 1 and 14 post T cell injection are shown. α-MOG CAR CD4+ T cells preferentially localized and expanded in the head area with a strong signal coming from the eyes compared to α-CD19 CAR CD4+ T cells control. n=3, mean +/- sem. **c**, Human α-MOG CAR CD4+ T cells lead to increased ventricle volume. NSG mice were injected with 5 x 10^6^ primary human CD4+ T cells expressing α-MOG CAR or control α-CD19 CAR. Around two weeks later, a T2 weighted MRI was done, and ventricle volumes were quantified. Representative images and quantifications are provided. Yellow arrows indicate the ventricle. n=3, mean +/- std. **d**, Human α-MOG CAR CD4+ T cells induce neuronal damage. The neuronal damage was assessed by the presence of neurofilament in the serum by ELISA. Mice injected with human α-MOG CAR CD4+ T cells had higher levels of neurofilament in the serum than mice injected with control α-CD19 CAR T cells. mean +/- sem. Mann-Whitney test. **e**, CNS specific α-MOG CAR T cells activation. α-MOG CAR T cells were isolated from the central nervous system (spine and brain) and spleen tissues two weeks after injection and stained for CD25 to evaluate T cell activation. α-MOG CAR T cells showed high levels of activation, specifically in the CNS compared to the spleen tissue. n=3, mean +/- std. Unpaired t-test. **f**, CNS specific α-MOG CAR T cells Th1 polarization. α-MOG CAR T cells were isolated from the central nervous system (spine and brain) and spleen tissues two weeks after injection and stained for IFN-**γ** and IL-17 to evaluate their polarization state. α-MOG CAR T cells showed strong polarization towards a Th1 phenotype, specifically in the CNS tissue compared to the spleen tissue, as shown by representative density plot.

In summary, α-MOG CAR CD4+ T cells traffic to and are activated in the CNS and induce an encephalitis associated with neuronal damage, with corresponding clinical signs of disability accumulation.

### Myelin-targeted CD4+ T cells produce consistent, brain-predominant, sharp demyelinating lesions at MS-typical sites, with perivascular architecture of human MS tissue

Given evidence of neurodegeneration reflected by increased ventricular volume on MRI and elevated NfL levels, we conducted a histopathological analysis of the brains and spinal cords of α-MOG CAR CD4+ T cell-injected NSG mice. This revealed discrete demyelinating lesions throughout the brain and optic nerve. To determine how closely these lesions recapitulate features of human disease, we compared their architecture directly with MS autopsy material. In MS, demyelinating lesions occur throughout the brain, including in periventricular and juxtacortical regions, a distribution recognized on clinical MRI and evident in an unstained coronal section of autopsy brain tissue, where plaques are visible as discrete pale areas of the white matter (Fig. 3a, left and middle).

**Fig. 3:**
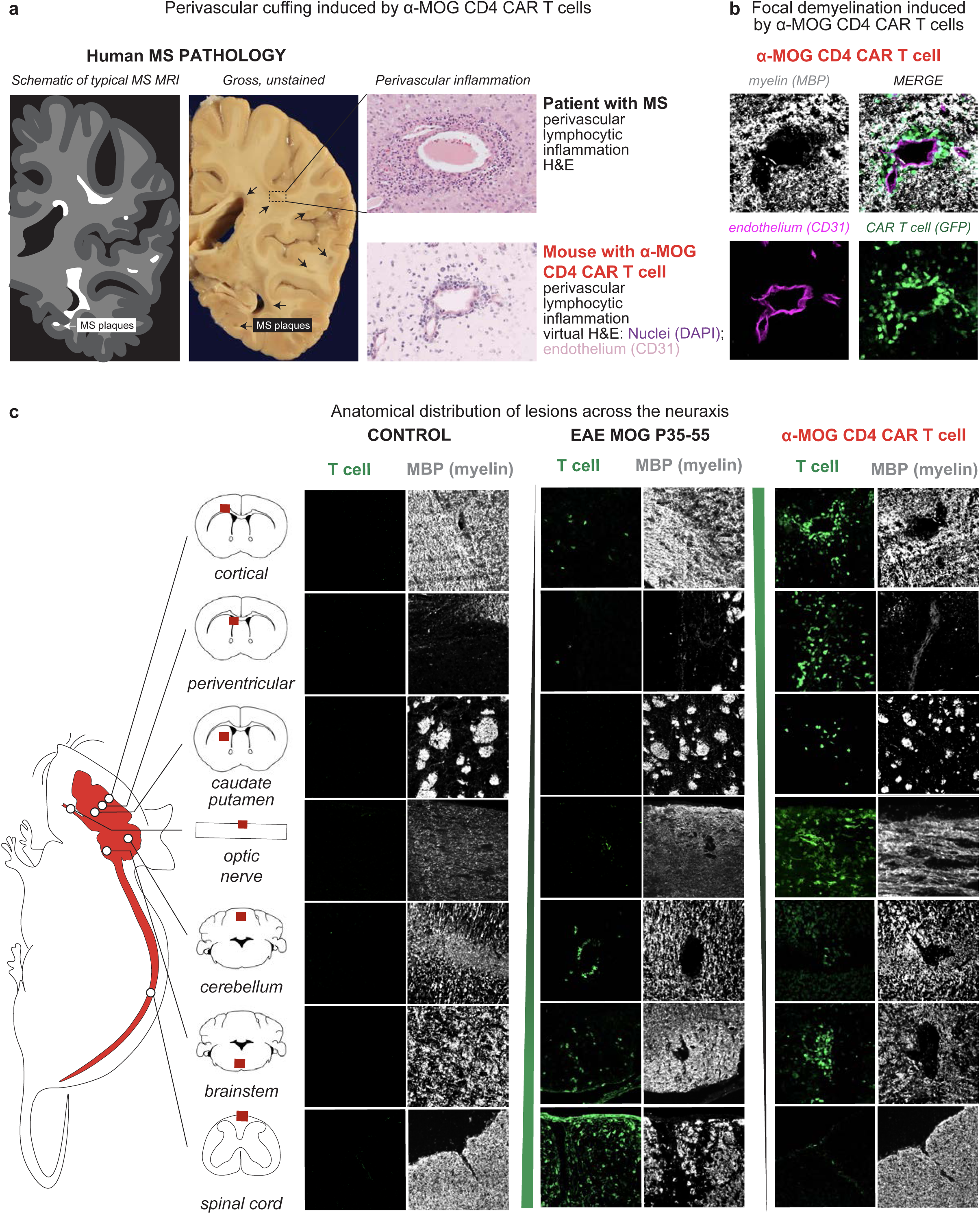
α-MOG CAR CD4+ T cells induce brain-predominant focal demyelination at MS-typical lesion sites, contrasting with the spinal cord-predominant diffuse pathology of EAE. **a**, Perivascular cuffing induced by α-MOG CAR T cells, compared with human MS pathology. Left, schematic illustration of the plaque distribution on a typical MS brain MRI corresponding to the middle, gross unstained coronal section of autopsy brain tissue from a patient with MS; arrows indicate demyelinating plaques. Right, perivascular lymphocytic infiltration in a different patient with MS (top, hematoxylin and eosin) and in a 5 x 10^6^ primary human α-MOG CAR CD4+ T cell-injected NSG mouse two weeks after adoptive transfer (bottom, virtual hematoxylin and eosin rendering of immunofluorescence for nuclei (DAPI) and endothelium (CD31); see Methods). In both species, mononuclear cells form a dense cuff around a small parenchymal vessel. **b**, Focal demyelination induced by α-MOG CAR T cells. The same cortical lesion as in **a**, was stained for MBP (myelin, grey) and CD31 (endothelium, magenta), with CAR T cells identified by constitutive GFP expression (green); merge, top right. CAR T cells accumulate around the CD31+ vessel at the center of a sharply demarcated MBP-negative lesion. Representative images. **c**, Anatomical distribution of lesions across the neuraxis. Cortical, periventricular, caudate putamen, optic nerve, cerebellum, brainstem and spinal cord were stained for MBP (myelin) and T cells (CD3 when no genetically encoded marker was used) in three groups: control mice (untreated / α-CD19 CAR CD4+ T cell-injected), C57BL/6J mice immunized with MOG p35-55 to induce EAE (day 17 post-immunization; T cells detected with an anti-CD3 antibody), and NSG mice injected with primary human CD4+ T cells expressing α-MOG CAR and GFP (two weeks after transfer; T cells detected by GFP). The schematic on the left indicates the sampled regions; the green gradient indicates regions most affected in each model. α-MOG CAR CD4+ T cells preferentially infiltrate anterior CNS regions - cortical, periventricular, caudate putamen, optic nerve, cerebellum, and brainstem - producing focal MBP loss at these sites and minimal spinal cord involvement. EAE mice show the reciprocal pattern, with spinal cord-predominant infiltration and diffuse rather than focal myelin loss, particularly evident in the spinal cord. Control mice show no T cell infiltration and intact MBP staining at all levels. Representative images. The same representative image of the cortical region as in **a** and **b** is shown.

Microscopically, MS plaques can contain a dense mononuclear cuff within the perivascular space, as shown by hematoxylin and eosin staining of autopsy tissue from a second case (Fig. 3a, top right). This perivascular organization is central to MS lesion formation, with the perivascular space acting as a privileged site for immune cells to interact with antigen-presenting cells, and the central vein sign that reflects this origin has recently been incorporated into the MS diagnostic criteria^18^. Lesions induced by α-MOG CAR CD4+ T cells reproduced this characteristic perivascular organization. To facilitate direct comparison with human histology, we rendered the mouse DAPI and CD31 immunofluorescence in hematoxylin and eosin-like contrast. The resulting images showed dense mononuclear infiltrate arranged concentrically around a parenchymal vessel and localized within the perivascular space (Fig. 3a, bottom right). Staining of the same cortical lesion for MBP and CD31 identified the CD31+ vessel at the center of a sharply demarcated MBP-negative area containing CAR T cells (Fig. 3b), directly linking the perivascular infiltrate to focal demyelination.

We next asked how these lesions were distributed across the neuraxis. We examined regions characteristically affected in MS, including the cortex, periventricular regions, optic nerve, cerebellum, brainstem, and spinal cord, in three different groups: control mice, C57BL/6J mice immunized with MOG p35-55 to induce EAE, and α-MOG CAR CD4+ T cell-injected NSG mice. Sections were stained for MBP and for infiltrating T cells (Fig. 3c). α-MOG CAR CD4+ T cells infiltrated every brain region examined as well as the optic nerve, producing focal MBP loss at each of these sites, while examination of the spinal cord revealed minimal or no lesions. This brain- and optic nerve-predominant distribution was consistent with the stronger bioluminescent signal in the head area than in the spinal cord (Fig. 2b), while involvement of the cerebellum and brainstem provides a plausible anatomical substrate for the ataxia observed in affected mice. Conventional EAE mice showed a strikingly different pattern: T-cell infiltration was spinal cord-predominant, and myelin loss was diffuse rather than organized in focal lesions, most evident in the spinal cord. Control mice showed no T cell infiltration and intact MBP staining at all levels examined (Fig. 3c).

Overall, these findings demonstrate that α-MOG CAR CD4+ T cells induce a distinct brain- and optic nerve-predominant demyelinating disease with limited spinal cord involvement. Importantly, the resulting focal lesions recapitulate two defining features of human MS pathology: their distribution across multiple clinically relevant CNS regions and their organization around a central parenchymal vessel. This pattern contrasts with the spinal cord-predominant, more diffuse inflammatory demyelination observed in conventional EAE, highlighting the complementary value of the CAR T cell model for studying focal CNS demyelination while preserving the established utility of EAE for investigating spinal cord inflammation and disability.

### Myelin-targeted CD4+ T cells cause reproducible severe optic neuritis with prominent discrete demyelinating lesions within the optic nerve

Another striking feature of this model is the consistent development of severe optic neuritis and discrete demyelinating lesions within the optic nerve. This was noted due to the notable bioluminescent signal from the orbital cavity (Fig. 2b), along with the marked bilateral periorbital alopecia (Fig. 4a), which suggested inflammation in the orbital cavity. Optic nerve involvement was already evident in the neuraxis survey described above, where the optic nerve was among the most heavily infiltrated sites (Fig. 3c). We, therefore, analyzed the retina and optic nerves via histology for possible optic neuritis. We observed a dramatic infiltration of α-MOG CAR CD4+ T cells by immunofluorescence histology in 100 % of the mice (9 out of 9 mice, across independent experiments, Extended Data Table 1), consistent with the bioluminescent signal. Additionally, we stained for and counted doublets of the paranodal contactin-associated protein (CASPR) that flank the nodes of Ranvier of the myelin sheath, which are essential for the ongoing depolarization of the axon and signal propagation^19^ (Fig. 4b). The α-MOG CAR CD4+ T cell-injected mice had a significant loss of paranodes, suggesting severe demyelination.

**Fig. 4:**
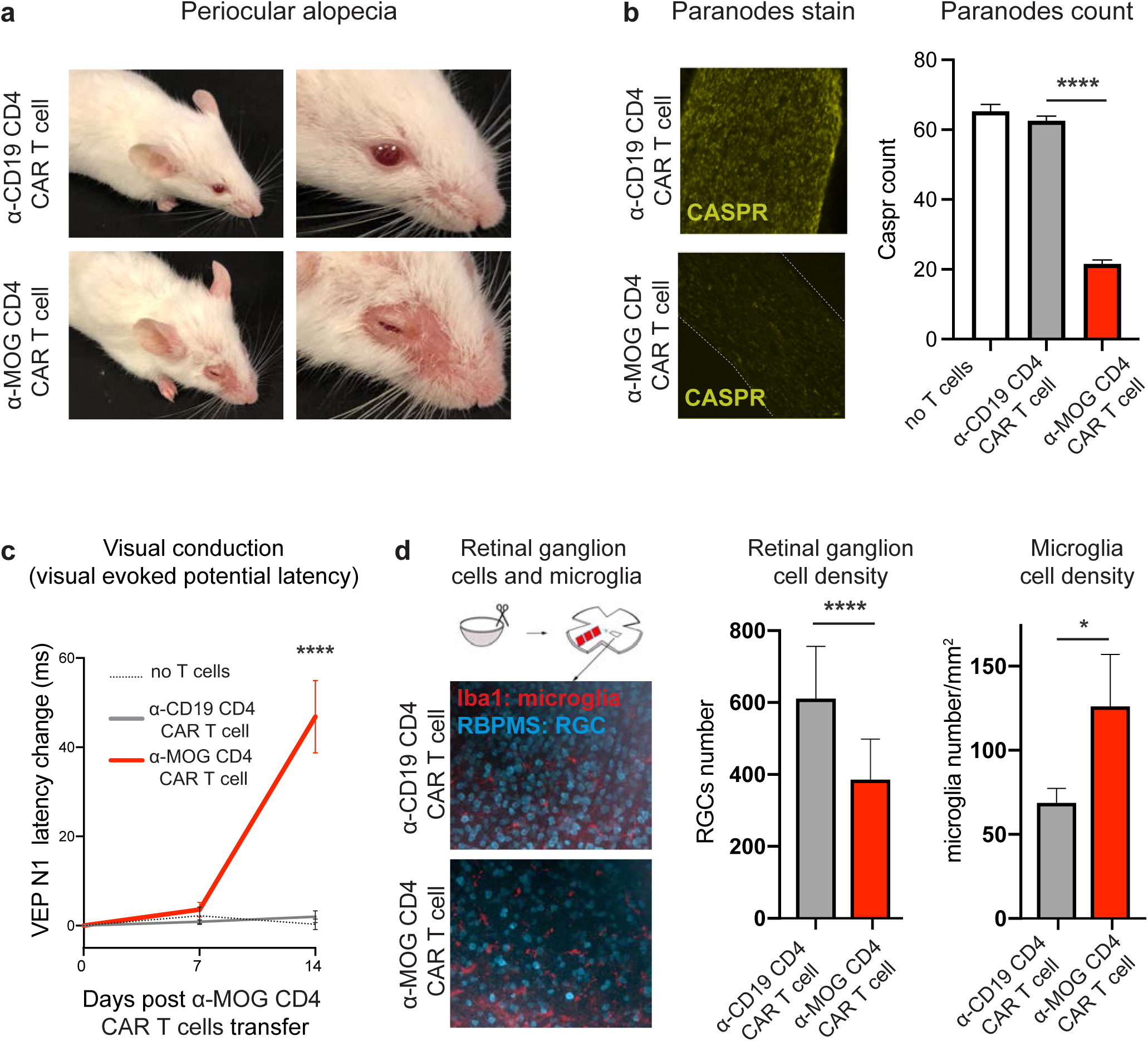
Human α-MOG CAR CD4+ T cells induce severe optic neuritis. **a**, Human α-MOG CAR CD4+ T cells affect the eye region of the mice. NSG mice were injected with 5 x 10^6^ primary human CD4+ T cells expressing α-MOG CAR or control α-CD19 CAR T cells. α-MOG CAR CD4+ T cells caused orbital bilateral alopecia and redness around the eye. Representative images (∼2-3 weeks after injection). **b**, Human α-MOG CAR CD4+ T cells demyelinate the optic nerves. The myelin state was assessed by CASPR stain, a paranode marker. Mice injected with human α-MOG CAR CD4+ T cells have far fewer paranodes than mice injected with control α-CD19 CAR T cells. Representative images and paranode count quantification. mean +/- std. Dunnett’s multiple comparison test. **c**, Human α-MOG CAR CD4+ T cells cause increased visual evoked potential latency. The visual evoked latency was measured over two weeks. Mice injected with α-MOG CAR CD4+ T cells had increased VEP latency consistent with demyelination shown in **b**. mean +/- std. **d**, Human α-MOG CAR CD4+ T cells cause retinal ganglion cell loss and microglial density increase. Retina flat mounts at day 14 after treatment with T cells were stained for RBPMS and Iba1 to respectively count the retinal ganglion cells and microglia. Mice injected with α-MOG CAR CD4+ T cells had fewer retinal ganglion cells and more microglia than mice injected with control α-CD19 CAR T cells. Representative images and retinal ganglion cell and microglia density quantification. mean +/- std. Unpaired t-test.

To functionally confirm optic nerve demyelination, we performed visual evoked potentials (VEP), an electrophysiological assay^20^. The VEP measures the electric potential generated by the visual cortex in response to a visual stimulus, enabling assessment of signal conduction time from the retina to the visual cortex^21,22^. Therefore, VEP measures are an objective readout of functional myelin status. The VEP latency increased only in the α-MOG CAR CD4+ T cell-injected mice (+45.6 ms, p < 0.01) (Fig. 4c), consistent with reduced signal conduction. Next, we assessed retinal ganglion cell loss, an established biomarker of neuroaxonal damage in MS patients with a history of optic neuritis^23^. Using retina flat mounts, we stained for the RNA-binding protein RBPMS, a marker of ganglion cells^24^, and Iba-1, a microglial marker. The density of retinal ganglion cells was markedly reduced in α-MOG CAR CD4+ T cell-injected mice, while the microglial cell density was increased (Fig. 4d). To assess the microglia state, we used an automated analysis to quantify the number of processes. The number of processes was decreased, suggesting a more amoeboid or activated state in α-MOG CAR CD4+ T cell-injected mice (Extended Data Fig. 4)^25^. When we looked for α-MOG CAR CD4+ T cells, we observed that CAR T cells were primarily localized in the optic nerve head and seemed to be distributed from there to the retina’s periphery (Extended Data Fig. 5). This is consistent with previous pathological studies describing retinal inflammatory cells in patients with chronic MS^26^.

Furthermore, when we stained the optic nerves with myelin basic protein (MBP), the staining was characterized by prominent discrete demyelinating lesions in the mice that received the α-MOG CAR CD4+ T cells (Fig. 5a), with the presence of α-MOG CAR CD4+ T cells (Extended Data Fig. 6). This was unlike the diffuse lesions observed in the traditional EAE immunization model using MOG P35-55 peptide (Fig. 5b).

**Fig. 5:**
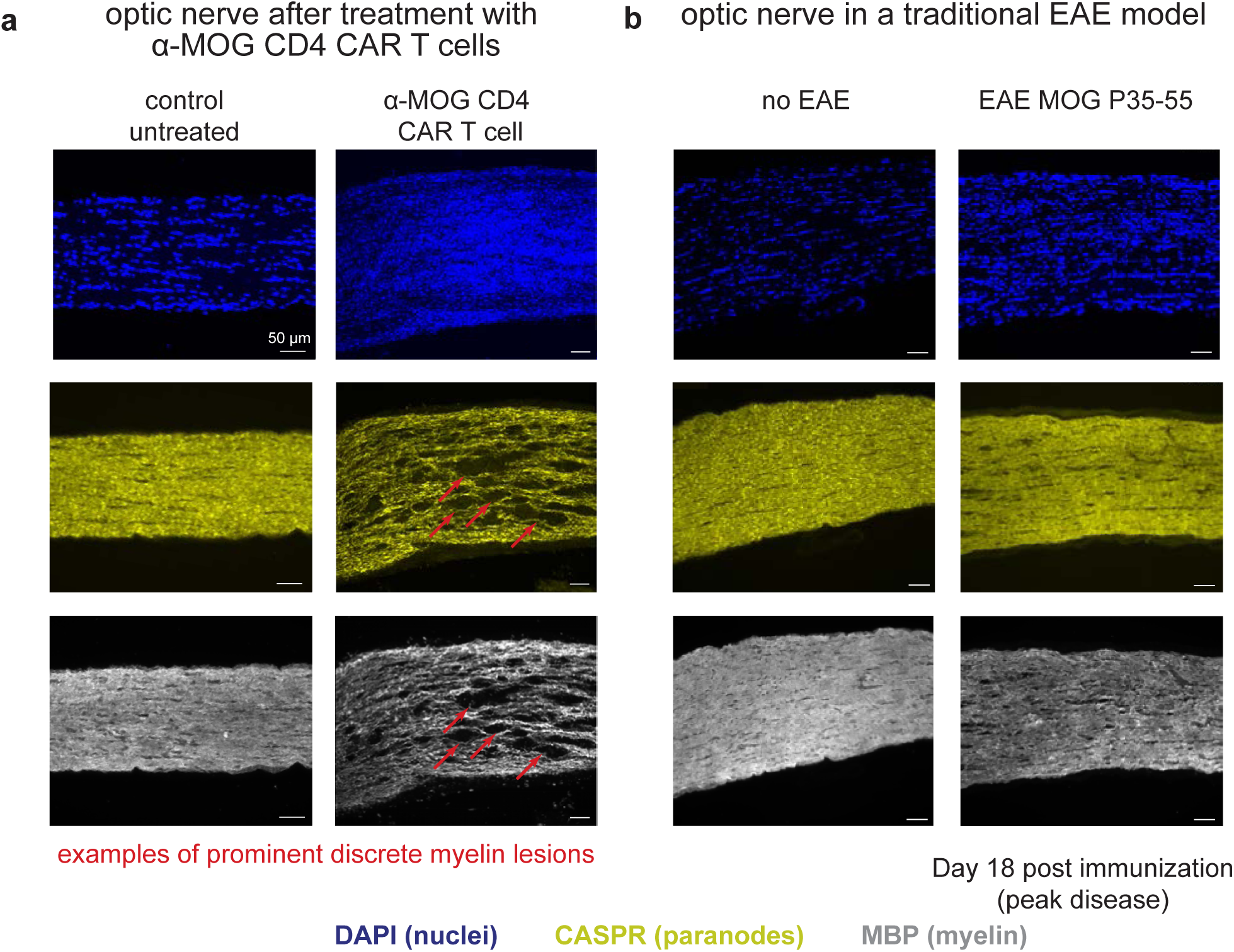
Human α-MOG CAR CD4+ T cells induce demyelination and prominent discrete lesions in the optic nerve resembling human disease (unlike EAE models). **a**, Human α-MOG CAR CD4+ T cells lead to prominent discrete demyelinating lesions. NSG mice were injected with 5 x 10^6^ primary human CD4+ T cells expressing α-MOG CAR. Two weeks later, the optic nerves were collected and stained for DAPI (nuclei), CASPR (paranodes), and MBP (myelin). α-MOG CAR CD4+ T cells lead to prominent discrete demyelination (red arrow), like in human optic neuritis. Control untreated NSG mice were collected and stained for DAPI (nuclei), CASPR (paranodes), and MBP (myelin). No lesions can be observed. **b**, Traditional EAE models do not show prominent discrete demyelinating lesions. C57BL/6J mice were immunized using a classical EAE model. The optic nerves were collected on day 18 post-immunization and stained for DAPI (nuclei), CASPR (paranodes), and MBP (myelin). Optic nerves show diffuse lesions at day 18 unlike human optic neuritis. Control non-immunized mouse (no EAE) is included for reference.

Altogether, these observations indicate that α-MOG CAR CD4+ T cells induce a reproducible, severe demyelinating optic neuritis characterized by prominent, discrete lesions; features absent in standard EAE. In contrast, MOG EAE mice exhibit diffuse disruption of nodal architecture with comparatively less pronounced focal loss of MBP staining.

### Clemastine, a remyelinating drug, restores optic nerve myelin and VEP latency in the CAR T-induced demyelination model

Given our model’s reliable optic neuritis features with prominent and easily identifiable discrete demyelinated lesions, we asked whether this model could be used to assess the efficacy of remyelinating drugs. The small molecule clemastine has undergone histological and electrophysiological in vivo validation following demyelination induced by EAE and cuprizone and has also been tested in clinical trials in MS^13,17,27,28^. Clemastine is a potent myelin regenerative compound that exerts its effect by antagonizing the muscarinic acetylcholine receptor 1 (M1R) on oligodendrocyte precursor cells^13^. Treatment with clemastine supports axonal integrity and neuronal function in inflammatory demyelination^13,17^. At day one after α-MOG CAR CD4+ T cell injection, mice were treated with clemastine or vehicle daily until day 14, similar to previous EAE trials^20^ (Fig. 6a). Mice treated with clemastine showed shorter VEP latency (Fig. 6b). Immunostaining of the optic nerves showed overall reduced volume of lesions as assessed by MBP and CASPR stains (Fig. 6c and Extended Data Fig. 7) (from 37% of the optic nerve area to 3.9%, p < 0.05). This was even more remarkable as the α-MOG CAR CD4+ T cells were still present throughout the optic nerve. To rule out that clemastine could inhibit α-MOG CAR CD4+ T cells, we cultured the T cells with K562 MOG^+^ cells over five days with various concentrations of the drug. We found that the range of in vivo clemastine concentrations had no detectable impact on α-MOG CAR CD4+ T cell activation or proinflammatory cytokine secretion (Extended Data Fig. 8). This result demonstrates that clemastine can promote reparative processes even during active inflammation in the presence of the α-MOG CAR T cells. Overall, α-MOG CAR CD4+ T cell-injected mice provide a robust model to assess the remyelinating effect of clemastine.

**Fig. 6:**
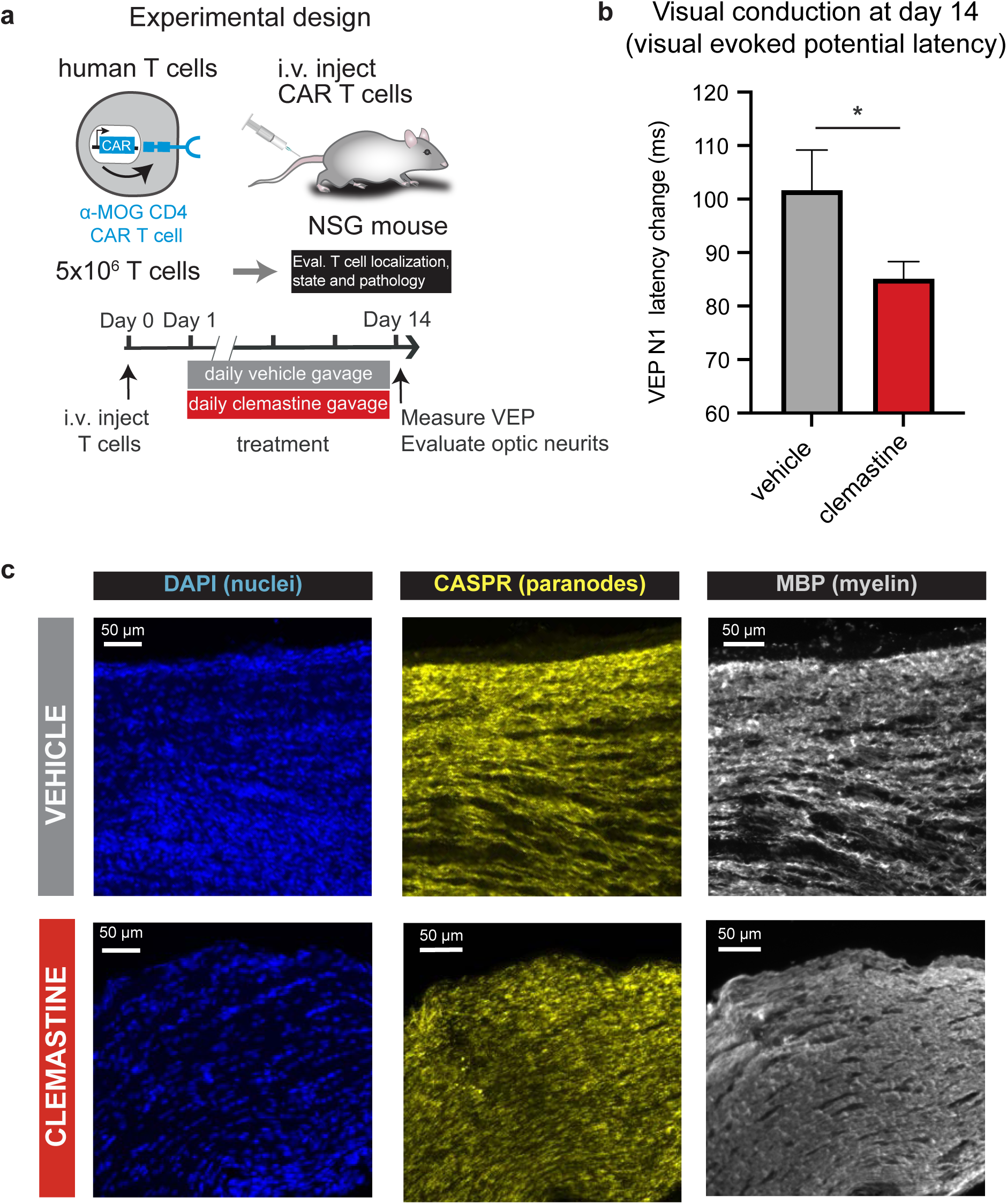
Clemastine treatment promotes remyelination and rescues α-MOG CAR CD4+ T cells induced severe optic neuritis. **a**, In vivo experimental design to assess clemastine treatment. Primary human CD4+ T cells were engineered to express α-MOG CAR and GFP-2A-effLuc. 5 x 10^6^ primary human CD4+ T cells were injected intravenously (i.v.) in immunocompromised NSG mice and treated with vehicle (n=6) or clemastine (n=8) by daily gavage starting at day one post T cell injection. Mice were followed for VEP and optic nerve staining evaluation. **b**, Clemastine decreases the VEP latency caused by the human α-MOG CAR CD4+ T cells. The visual evoked latency was measured at day 14 after T cell injections. Mice treated with clemastine had lower VEP latency. mean +/- sem. Unpaired t-test. **c**, Clemastine protects from the demyelination caused by the human α-MOG CAR CD4+ T cells. The optic nerves were processed by immunofluorescence and stained for DAPI (nucleus), CASPR (paranodes) and MBP (myelin). Clemastine-treated optic nerves had more intact myelin despite having α-MOG CAR T cells present (Extended Data Fig. 7).

## DISCUSSION

Animal models with reproducible disease and the fidelity to recapitulate discrete demyelinating lesions seen in people with MS are greatly needed in the field of demyelinating diseases to help overcome the barrier to remyelination. In this work, we have leveraged the CAR system, by engineering primary human CD4+ T cells to target the myelin protein MOG, to create a reproducible disease process of brain-directed inflammatory demyelination. These cells enter the CNS and optic nerve, where they are activated, generating a lesional demyelinating pathology resulting in ongoing clinical disability accumulation and neuronal loss; features not observed in traditional EAE models.

This work establishes a broader conceptual framework for disease modeling where engineered cells are used as precise in vivo perturbative agents to isolate and study specific pathological features of complex inflammatory diseases, independent of their underlying causality. Rather than replicating the full complexity of human MS immunopathogenesis, this approach reliably generates critical disease features, such as discrete demyelinating lesions in anatomically consistent areas, that enable targeted interrogation of lesion predilection, remyelination biology, and therapeutic intervention. Our model demonstrates this principle through its striking anatomical similarity to MS, with a brain-predominant lesion distribution spanning regions central to MS pathology (juxtacortical, periventricular, cerebellar, brainstem, and optic nerve) and perivascular infiltration consistent with the central vein sign. Side-by-side comparison with MS autopsy tissue shows that this correspondence holds at the level of both lesion topography and perivascular architecture (Fig. 3a). Notably, both the optic nerve and the central vein sign were formally incorporated into the 2024 McDonald diagnostic criteria for MS^18^.

In this study, we used the well-established autoantigen MOG as a CAR target. The fidelity and reproducibility of this approach, however, suggest the potential to engineer additional antigen-specific disease models across a range of disorders, including paraneoplastic neurological disorders^29^. More broadly, CARs could be introduced into diverse immune cell types: CD8+ T cells, CD4+ Th1 or Th17 T cells, NK cells, or macrophages/microglia, to investigate how different effector cells targeting the same antigen influence disease and delineate their respective contributions.

The pathology and histology of demyelination in EAE models vary based on the strain and the method used for induction: 1) immunization vs 2) adoptive transfer or 3) spontaneous EAE^30^. A discussion about the different models’ pathology is beyond the scope of this work, but in the most frequently used model of EAE, C57BL/6 mice immunized with MOG35–55 peptide, the spinal cord exhibits multifocal, confluent demyelination in the peripheral white matter; the brain shows meningitis with perivascular inflammatory cuffing in the cerebellum and hindbrain white matter (Fig. 3c); and the optic nerves display generalized demyelination (Fig. 5). However, in that model there is no CNS region discrete lesions similar to the neuropathology in human demyelinating disorders such as MS^30^. For this reason, the field has been unable to investigate whether remyelinating compounds act on oligodendrocyte progenitor cell differentiation only in the context of normal-appearing white matter or within lesions. Data from imaging biomarkers from two different clinical trials, ReBUILD^28^ and CCMR One^31^, evaluating the safety of two different remyelinating compounds (clemastine and bexarotene), have not answered this question with certainty. Our new model with discrete demyelinating lesions overcomes these neuropathological differences between EAE and MS and shows that clemastine can induce lesion remyelination in the acute inflammatory phase of demyelination. This also highlights the value of this approach as a potential preclinical testbed for the assessment of new remyelinating therapies. Furthermore, besides lesions, this model allows for exploring additional aspects of optic neuritis, including the unexplained infiltration of inflammatory cells targeting myelin through the optic nerve head into retinal layers (Extended Data Fig. 5). A shortcoming of this model is the limited spinal cord involvement; EAE thus remains the preferred model for studying spinal cord pathology, and the two approaches are best viewed as complementary tools for addressing the spectrum of MS pathology.

Our model shows that a monospecific T cell population is sufficient to initiate demyelinating disease in an immunocompromised mouse. This finding has significant implications for understanding the factors contributing to disease development, suggesting that focal lesion formation may not require complex polyclonal responses or a full immune system response. While this model reliably induced disease, we noticed some variations in the disease intensities depending on the donor of the primary human T cells that could be explored in further studies.

In conclusion, the fidelity, reproducibility, and human-like pathophysiology make this model advantageous for studying CNS demyelination disease. As a proof-of-principle, it highlights the potential of engineered cells to open new avenues for clinical translation, advancing remyelination research, and enabling the modeling of a broader spectrum of inflammatory disorders.

## METHODS

### Construct Design

CARs were built by fusing CD8a signal peptide (MALPVTALLLPLALLLHAARP) for membrane targeting and α-myc-tag (EQKLISEEDL) for detecting surface expression with α-myc A647 (cell-signaling #2233) to the α-MOG clone M26^32^ or α-CD19 clone FMC63^33^ binders to the hinge region of the human CD8α chain and transmembrane and cytoplasmic regions of the human 4-1BB, and CD3z signaling domains. Receptors were cloned into a modified pHR’SIN:CSW vector containing a PGK promoter.

### Primary Human T Cell Isolation and Culture

Primary CD4+ T cells were isolated from donor blood after apheresis by negative selection (STEMCELL Technologies). Blood was obtained from StemExpress or Allcells, as approved by the University of California San Francisco (UCSF) Institutional Review Board. T cells were cryopreserved either in Cellbanker 1 or in RPMI-1640 with 20% human AB serum (Valley Biomedical) and 10% dimethyl sulfoxide. After thawing, T cells were cultured in human T cell medium consisting of X-VIVO 15 (Lonza), 5% Human AB serum, 55 µM b-mercaptoethanol and 10 mM neutralized N-acetyl L-Cysteine supplemented with 30 units/mL interleukin (IL)-2.

### Lentiviral Transduction of Human T Cells

Pantropic vesicular stomatitis virus G (VSV-G) pseudotyped lentivirus was produced via transfection of Lenti-X 293T cells (Takara Bio, Cat# 632180) with a pHR’SIN:CSW transgene expression vector and the viral packaging plasmids pCMV and pMD2.G using Fugene HD (Promega) or TransIT-VirusGen (Mirus Bio). Primary T cells were thawed the same day and, after 24 hours in culture, were stimulated with 25 µl anti-CD3/CD28 coated beads [Dynabeads Human T-Activator CD3/CD28 (Gibco)] per 1 ×10^6^ T cells. In some instances, freshly isolated T cells were used. At 48 hours, viral supernatant was harvested. Primary T cells were exposed to the lentivirus for 24 hours or spinoculated on retronectin-coated plates. At day 5 after T cell stimulation, Dynabeads were removed, and T cells were stained with an anti-myc tag AF647 antibody (Cell Signaling 2233) and sorted with a Becton Dickinson (BD) FACS ARIA Fusion or Sony SH800S Cell Sorter. T cells were expanded until rested and could be used in assays.

### Human T Cell Staining and Phenotyping

T cell phenotypes were assessed using the following antibodies: Fluorescein isothiocyanate (FITC) anti-CD3 (clone HIT3a, 300306, Biolegend), allophycocyanin (APC) anti-CD25 (clone BC96, 302610, BioLegend) for T cell activation; PE-Cy7 anti-PD1 (clone EH12.1, 561272, BD Biosciences), BV785 anti-TIM3 (clone F38–2E2, 345031, BioLegend), and AF700 anti-LAG3 (clone 3DS223H, 56–2239-42, Thermo Fisher Scientific) for T cell exhaustion; BV605 anti-CD45RA (clone HI100, 304133, BioLegend) and allophycocyanin (APC)–Cy7 anti-CD62L (clone DREG-56, 304813, BioLegend) T cell differentiation state. For intracellular stains, after 4 hours of incubation with GolgiPlug^TM^ (555029, BD Biosciences) and GolgiStop^TM^ (554724, BD Biosciences), we used: PE anti-IL17A (clone BL168, 512305, BioLegend) and APC anti-IFN-γ (clone 4S.B3, 502512, BioLegend).

Cells were analyzed by flow cytometry using BD Fortessa; analysis was performed by FlowJo software (TreeStar).

### Cell Lines

Cell lines used were K562 myelogenous leukemia cells (ATCC #CCL-243). Cells were lentivirally transduced to stably express mCherry, and enhanced firefly luciferase under control of the spleen focus-forming virus (SFFV) promoter and sorted as previously shown^34^. For transgene expression, K562s were transduced with the lentiviral vector CD510-B1 (System Biosciences) and puromycin selected. K562 cells were cultured in DMEM 10% FBS.

### In Vitro Stimulation of α-MOG CAR T cells

For in vitro stimulation, the engineered α-MOG CAR T cell and target K562 cells were co-cultured at a 1:1 ratio, 1 x 10^5^ cells each in a flat bottom 96-well tissue culture plate for the indicated times and stained accordingly to evaluate activation, proliferation, differentiation, and exhaustion. When cytokine release assays were conducted, the supernatant was collected and processed by ELISA (IFN-γ, 430104, IL-6, 430504, and TNF-α, 430204, BioLegend).

### Assessment of α-MOG CAR T Cell Cytotoxicity

CD4+ α-MOG CAR T cells or control non-transduced T cells were cultured for 72 hours with target K562 MOG^+^ cells (1:1, 1 × 10^4^). The degree of specific lysis of target cells was determined by comparing the fraction of target cells alive in the culture compared to treatment with non-transduced T cell controls.

### In Vivo Mouse Experiments

All mouse experiments were conducted according to Institutional Animal Care and Use Committee (IACUC)–approved protocols.

8- to 12-week-old female NSG mice initially purchased from Jackson laboratory were injected tail-vein with 5 x 10^6^ α-MOG or α-CD19 CAR T cells in 100 μL. Survival was evaluated over time until predetermined IACUC-approved endpoint (hunching, neurological impairments such as circling, ataxia, paralysis, limping, head tilt, balance problems, seizures, body-weight loss) was reached.

Longitudinal T cell localization analysis was evaluated by luminescence emission on Xenogen IVIS Spectrum after intraperitoneal injection of 1.5mg D-luciferin (GoldBio, injection volume 200 μl).

In the EAE experiments, wild-type female C57BL/6 (B6) mice (8-12 weeks of age) were purchased from Jackson Laboratory. Mouse MOG p35-55 (MEVGWYRSPFSRVVHLYRNGK) was purchased from Genemed Synthesis (San Antonio, TX). For Fig. 3, EAE was induced using the Hook Labs kit (EK2210). For Fig. 5, mice were immunized with 100 µg of MOG p35-55 emulsified in M. tuberculosis H37Ra (Difco Laboratories, Franklin Lakes, NJ) and incomplete Freund adjuvant (BD) subcutaneously (sc). Afterward, 200 ng of B. pertussis toxin (List Biological, Campbell, CA) was injected by the intraperitoneal (IP) route on the day of induction and 2 days later. All mice were monitored daily for signs of clinical EAE for 18 days.

### Quantification of NfL from the serum

Blood was collected from intracardial puncture and let to clot at room temperature followed by centrifugation to collect the serum. Sera were analyzed by an HD-X analyzer using an Nf-Light Advantage kit from Quanterix Corporation, Lexington, MA, USA. All samples were measured in replicates.

### T2 weighted MRI

All in vivo MR experiments were conducted on a 14.1-T vertical MR system (Agilent Technologies) equipped with 100 G/cm gradients and a 1H volume coil (inner diameter, 40 mm). For each imaging session, the mice were anesthetized using isoflurane (1.5%–2% in O2). The animals were placed in a water-heated cradle with the head secured to ensure similar positioning between experiments. Respiration was continuously monitored using PC-SAM software (SA Instruments, Inc.). T2-weighted MRI was performed using the following parameters: an echo time/repetition time of 11.8/3,000 ms, a rapid-imaging-with-refocused-echoes factor of 8, a slice thickness of 0.5 mm, a gap of 0.25 mm, 6 averages, a matrix of 256 × 256, and a field of view of 30 × 30 mm. For MR analyses the tissue surrounding the brain was masked and the volume of the ventricles (lateral and 3rd ventricles) was manually delineated using ITK-SNAP (version 3.8.0)^35^.

### Isolation of CNS Infiltrating Mononuclear Cells and Splenocytes

Mice were treated with 1 mL of Avertin and perfused using PBS; CNS tissue was isolated and cut into small pieces in RPMI. After centrifugation, the tissue was resuspended in 5mL of RPMI with DNAse I at 200 U/mL, Roche #04536282001) and collagenase D (2.5mg/mL, Roche #11088882001) and incubated for 20 min at 37C with occasional shake. The digested sample was strained through a 70 µm cell strainer. After centrifugation, the pellet was resuspended in 6mL of low-density Percoll solution (1.030, consisting of 1.5mL Percoll + 5mL completed T cell media containing 10% FBS) and overlayed onto 6mL of high-density Percoll (1.095, consisting of 5mL Percoll + 1.84mL PBS). Centrifuge 15 min at 760g at room temperature with low brake. After removing the myelin debris, collect the mononuclear cells at the interphase. Wash with 50mL RPMI, centrifuge, and resuspend in the appropriate buffer. The spleens were strained through a 70 µm cell strainer and incubated with RBC lysis before washing and resuspension.

Cell surface staining was performed immediately, while intracellular stain was done after culturing the cells in GolgiPlug^TM^ (555029, BD Biosciences), GolgiStop^TM^ (554724, BD Biosciences), and PMA/ionomycin for 6 hours at 37C. Cells were then washed and fixed with Cytofix/Cytoperm™ (554714, BD Biosciences) followed by Perm/Wash™ (554723, BD Biosciences) and incubation with the appropriate antibodies.

### Immunofluorescence

Mice were treated with 1 mL of Avertin and perfused using PBS transcardially followed by PFA. Brains, spinal cords, and optic nerves were removed and fixed overnight in 4% paraformaldehyde–PBS before being transferred to 30% sucrose and allowed to sink (1-2 days). The eyes were transferred to PBS instead.

The optic nerves were embedded in Tissue-Tek OCT (Sakura) and long-term stored at −80°C. Immunohistochemistry was performed on 16 µm longitudinal sections. Sections were rinsed in PBS and blocked with 10% neutral goat serum and 0.05% Triton-X. Primary antibody incubations were carried out overnight at 4°C in blocking solution. The staining was performed using rabbit anti-CASPR (1:2000, Abcam, ab34151) and rat anti-MBP (1:200, Sigma-Aldrich, MAB386) as primary antibodies. Gt-αRb (1:1000)-647 and Gt-αRt (1:1000)-488 were used as secondary antibodies. Alexa-Fluor tagged secondary antibodies (Invitrogen) were used for primary antibody detection at room temperature. DAPI containing mounting media (Vectashield) was used to mount tissue to slide. Images were acquired using an upright Axio Imager 2 microscope (Carl Zeiss Microscopy) and processed using ImageJ software (NIH).

For flat mounted retinas, eyes were post-fixed in 4% PFA for one hour then moved to PBS. Retinas were dissected out of the eyes and washed with PBS before being placed in blocking buffer solution made of 10% BSA and 0.03% Triton-X in PBS. Primary antibody incubations were carried out overnight 4°C in blocking solution. Guinea pig anti-RBPMS (1:200, ThermoFisher, PA5-31231) and rabbit anti-Iba1 (1:1000, abcam, ab283346) were used for primary antibody staining. Tissue was then left to incubate in the dark for one hour using Gt-αGp (1:1000)-555 and Gt-αRb (1:1000)-647 as secondary antibodies. Retinas were transferred to slide and carefully cut as to create the clover shape, allowing the retinas to lay flat on the slide. Lastly, DAPI containing mounting media (Vectashield) was used to mount tissue.

### Human MS Autopsy Tissue and Neuropathology

Images of autopsy brain tissue from two patients with MS were used (Fig. 3a) and obtained from L.N.K. at the University of Chicago pathology department. From the first case, a coronal slice was photographed unstained to document the gross distribution of demyelinated plaques. From the second case, a formalin-fixed and paraffin-embedded block section containing a lesion was stained with hematoxylin and eosin. The accompanying MRI panel is a schematic illustration of the plaque distribution that corresponds to the gross specimen; note that it does not reproduce the actual patient imaging data.

### Virtual Hematoxylin and Eosin Rendering of Immunofluorescence Images

To allow direct visual comparison between mouse immunofluorescence and human hematoxylin and eosin histology (Fig. 3a), the DAPI (nuclei) and CD31 (endothelium) channels were pseudocolored in ImageJ/Fiji (NIH) to approximate hematoxylin and eosin contrast: the nuclear channel was assigned a hematoxylin-like blue-violet color and the endothelial channel an eosin-like pink color, both channels were inverted onto a white background, and the two channels were merged. Only look-up table (color) assignment, inversion, and linear brightness and contrast adjustments applied uniformly to the entire image were used; no local adjustment, filtering, or other structural manipulation was performed. Unrendered immunofluorescence source images are available from the corresponding authors on request.

### Visual evoked potentials (VEP) measurements

The VEP was measured as previously described ^20,36–38^. Briefly, the mice were anesthetized via intraperitoneal injection using a combination of xylazine (20 mg/ml) and ketamine (100 mg/ml) diluted in PBS administered at a dose of 0.125 ml per 20 g of mouse weight and positioned on a flat surface to insert the 1 cm steel needle electrodes (Natus Neurology). The active electrode for the flash VEP is placed medially under the skin between the two eyes, along the sagittal suture, with the needle’s tip inserted to 8 mm depth to optimize proximity to the visual cortex. A subcutaneous needle electrode inserted just above the nose tip serves as a reference, and a needle electrode in the tail serves as the ground. Each exam consisted of three runs, with pulse intensity 3 cd.s/m^2^, frequency 1 Hz, on-time 4 ms, white pulse color at 6500 K, and 100 sweeps per acquisition.

### Statistical Analysis

All statistical analyses were performed with Prism software version 9.0 (GraphPad) as described in the figures and legends.

## Data availability

All data are available in the manuscript or supplementary materials. Reagents are available from the corresponding author upon reasonable request. Plasmids from this paper will be made available on Addgene.

## Acknowledgments

We thank N. Blizzard, M. Broeker, P. Lopez Pazmino, S. Sidhu, and members of the W.A.L., S.S.Z., M.R.W. and M.S.S laboratories, as well as Dr. Peter Pytel for assistance, advice, and helpful discussions.

## Author contributions

Conceptualization: M.S.S., S.G., C.C., S.S.Z., A.J.G., and W.A.L.

Methodology: M.S.S., S.G., C.C., S.S.Z., A.J.G., and W.A.L.

Investigation: M.S.S., S.G., C.C., C.S., M.H., J.D., A.A., B.N., R.Z., W.Y., S.A.S., D. G. S., J.H.S., A.B., A.M., C.G., L.K., S.L.H., M.R.W., S.S.Z., A.J.G., and W.A.L.

Visualization: M.S.S., S.G., C.C., S.S.Z., A.J.G., and W.A.L.

Funding acquisition: M.S.S., S.G., C.C., M.R.W., S.S.Z., A.J.G., and W.A.L.

Project administration: M.S.S., M.R.W., S.S.Z., A.J.G., and W.A.L.

Supervision: M.S.S., M.R.W., S.S.Z., A.J.G, and W.A.L.

Writing original draft: M.S.S., S.G., C.C.

Writing review & editing: M.S.S., S.G., C.C., C. G., M.R.W., S.S.Z., A.J.G., and W.A.L.

## Ethics declarations

W.A.L. is shareholder of Gilead Sciences, Allogene, and Intellia Therapeutics, and previously consulted for Cell Design Labs, Gilead, Allogene, and SciFi Foods. S. L. H. currently serves on the scientific advisory board of Accure, Alector, Annexon. He has previously consulted for BD, Moderna, NGM Bio, Pheno Therapeutics and previously served on the Board of Directors of Neurona. Dr. Hauser also has received travel reimbursement and writing support from F. Hoffmann-La Roche and Novartis AG for anti-CD20-therapy-related meetings and presentations. M.R.W. is a co-founder and board member of Delve Bio, Inc., has received unrelated research grant funding from Roche/Genentech, Novartis, and Kyverna Therapeutics, and has received consulting fees from Pfizer, Indapta Therapeutics, Red Tree Ventures, Ouro Medicines, and Vertex Pharmaceuticals.

