## Extended Data for "Modeling CNS inflammatory demyelination using myelin-directed CAR T cells"

**Extended Data Fig. 1: Human α-MOG CAR CD4+ T cells polarize towards a Th1 response when engaging with MOG-expressing cells (related to Fig. 1)**

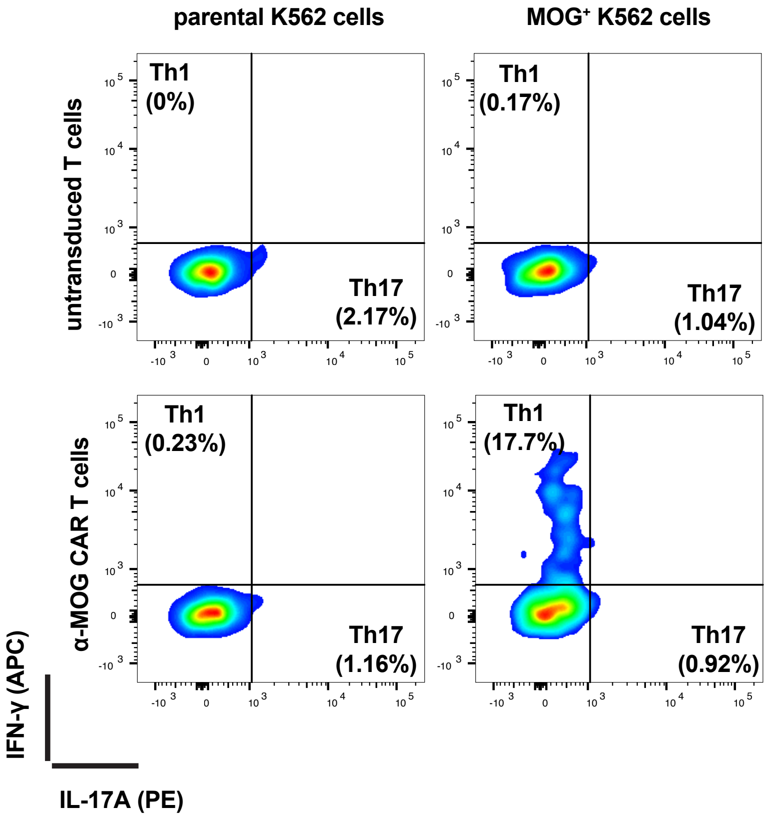


*E:T 1:1 10,000 cells*
*24h*

**Extended Data Fig. 1: Human α-MOG CAR CD4+ T cells polarize towards a Th1 response when engaging with MOG-expressing cells (related to Fig. 1).**

Th1 and Th17 polarization assessment in vitro. Primary human CD4+ T cells transduced with α-MOG CAR were co-cultured with K562 priming cells, either expressing or not expressing MOG, at indicated E:T ratio, duration, and cell numbers. Th1 and Th17 polarization was assessed by flow cytometry using an IFN-γ and IL-17A intracellular stain, respectively. Representative density plots are shown. n = 3.

**Extended Data Fig. 2: Human α-MOG CAR CD4+ T cells differentiate towards an effector memory state and upregulate exhaustion markers when engaging with MOG-expressing cells (related to Fig. 1)**

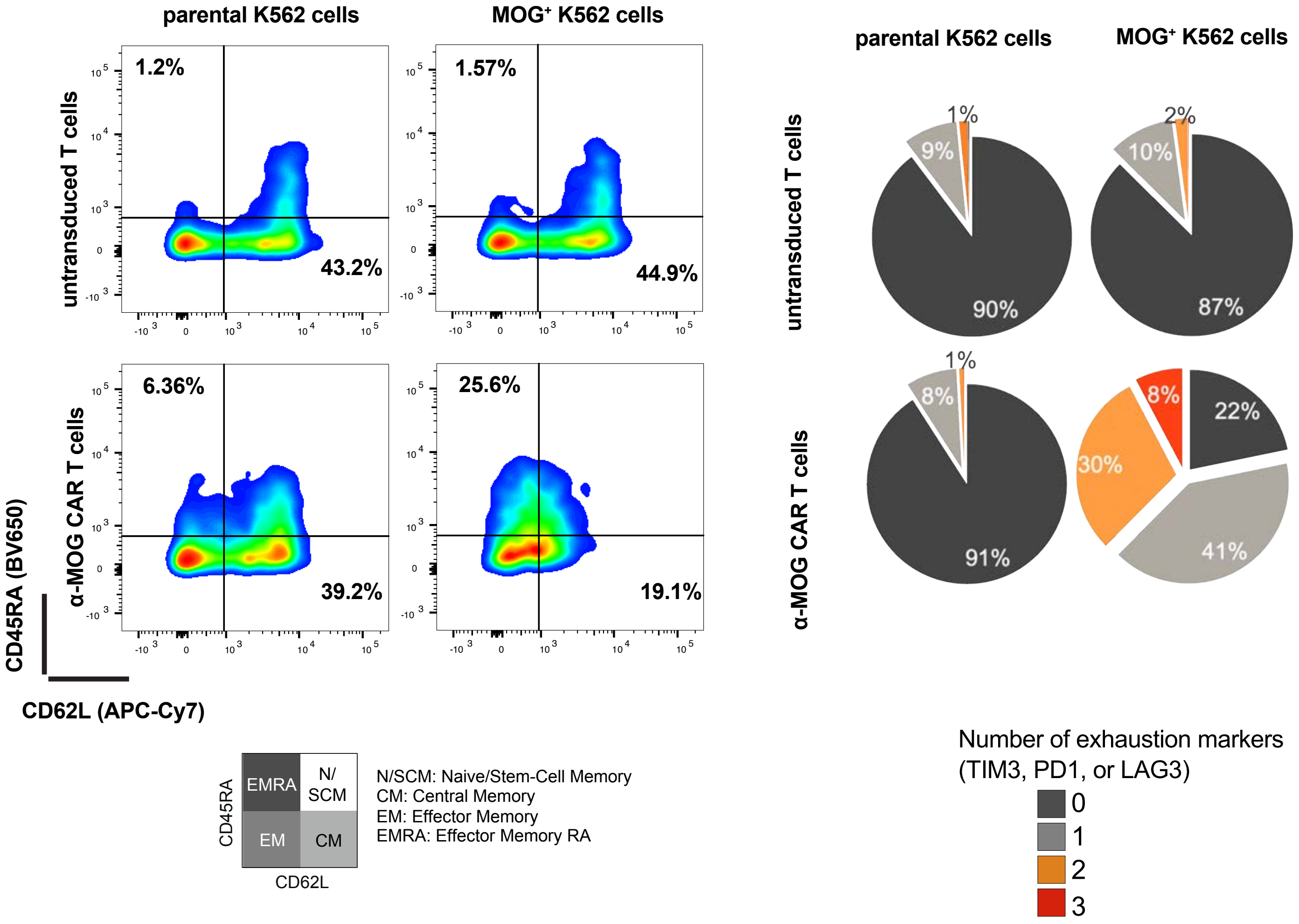


*E:T 1:1 10,000 cells*
*24h*

**Extended Data Fig. 2: Human α-MOG CAR CD4+ T cells differentiate towards an effector memory state and upregulate exhaustion markers when engaging with MOG-expressing cells (related to Fig. 1).**
T cell differentiation and exhaustion state assessment in vitro. Primary human CD4+ T cells transduced with α-MOG CAR were co-cultured with K562 priming cells, either expressing or not expressing MOG, at indicated E:T ratio, duration, and cell numbers. Flow cytometry plots distinguishing naïve-like cells ($CD45RA^+CD62L^+$), central memory cells ($CD45RA^-CD62L^+$), effector memory cells ($CD45RA^-CD62L^-$), and effector memory RA cells ($CD45RA^+CD62L^-$). Pie charts show the percentage of cells that express 0, 1, 2, or 3 exhaustion markers (PD1, LAG3, or TIM3) without and with stimulation by target cells, an average of three repeats.

**Extended Data Fig. 3: Human α-MOG CAR CD4+ T cell-injected mice show rapid body weight loss and death (related to Fig. 2)**

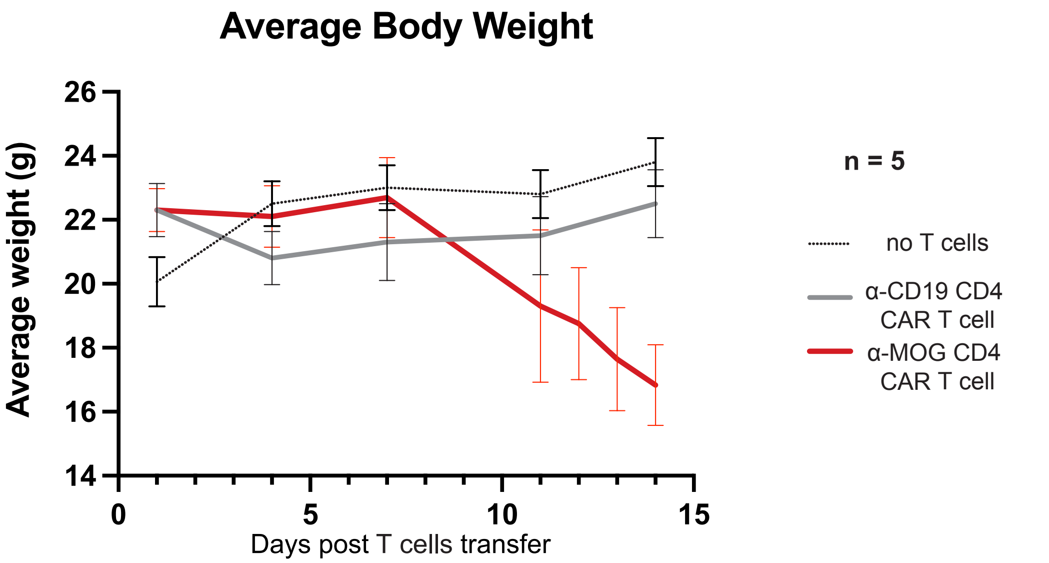


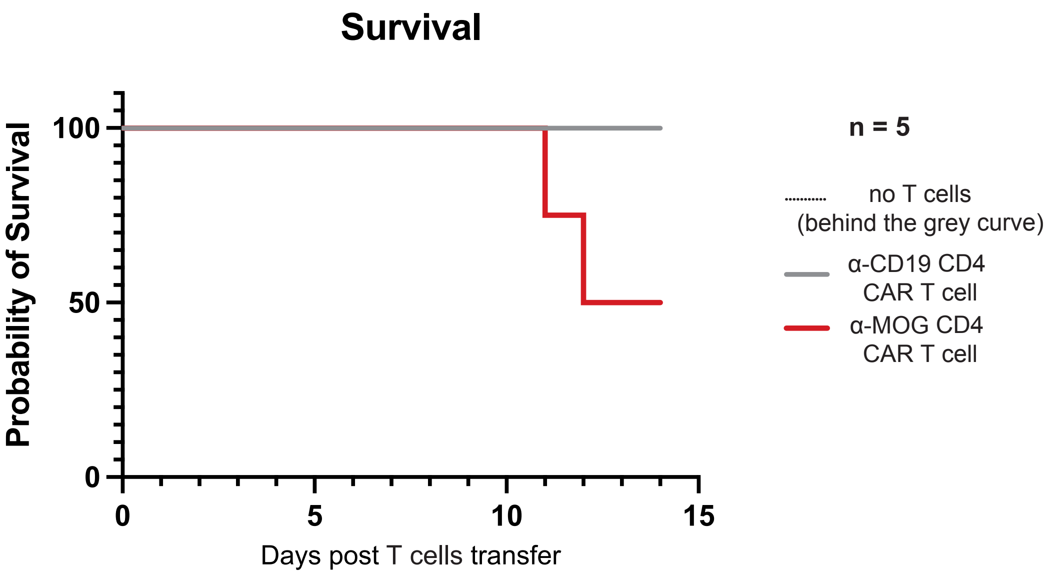


**Extended Data Fig. 3: Human α-MOG CAR CD4+ T cell-injected mice show rapid body weight loss and death (related to Fig. 2).**
NSG mice were injected with 5 x $10^6$ primary human CD4+ T cells expressing α-MOG CAR or control α-CD19 CAR. Body weight and survival were assessed for 14 days post adoptive transfer. n = 5 mice per group.

**Extended Data Fig. 4: Human α-MOG CAR CD4+ T cells induce microglia activation of the optic nerve (related to Fig. 4)**

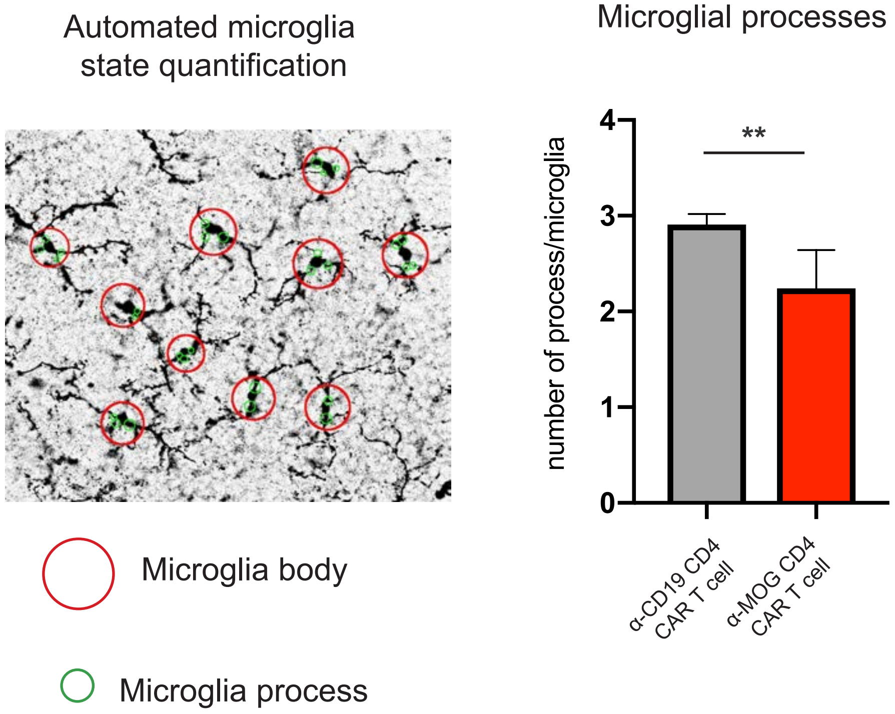


**Extended Data Fig. 4: Human α-MOG CAR CD4+ T cells induce microglia activation of the optic nerve (related to Fig. 4).**
Human α-MOG CAR CD4+ T cells affect the microglia state of the optic nerves. NSG mice were injected with 5 x $10^6$ primary human CD4+ T cells expressing α-MOG CAR or control α-CD19 CAR. Two weeks later, the optic nerves were collected and stained for Iba1 (microglial marker). α-MOG CAR CD4+ T cells caused changes in the microglia of the optic nerve. An automated script allows to identify microglia (red circle around the cell body) and the number of processes (green circles stemming from the cell body). Representative image.
Quantification of the number of processes per microglia is displayed. mean +/- std. Unpaired t-test.

**Extended Data Fig. 5: Human α-MOG CAR CD4+ T cells infiltrate the retina (related to Fig. 4)**

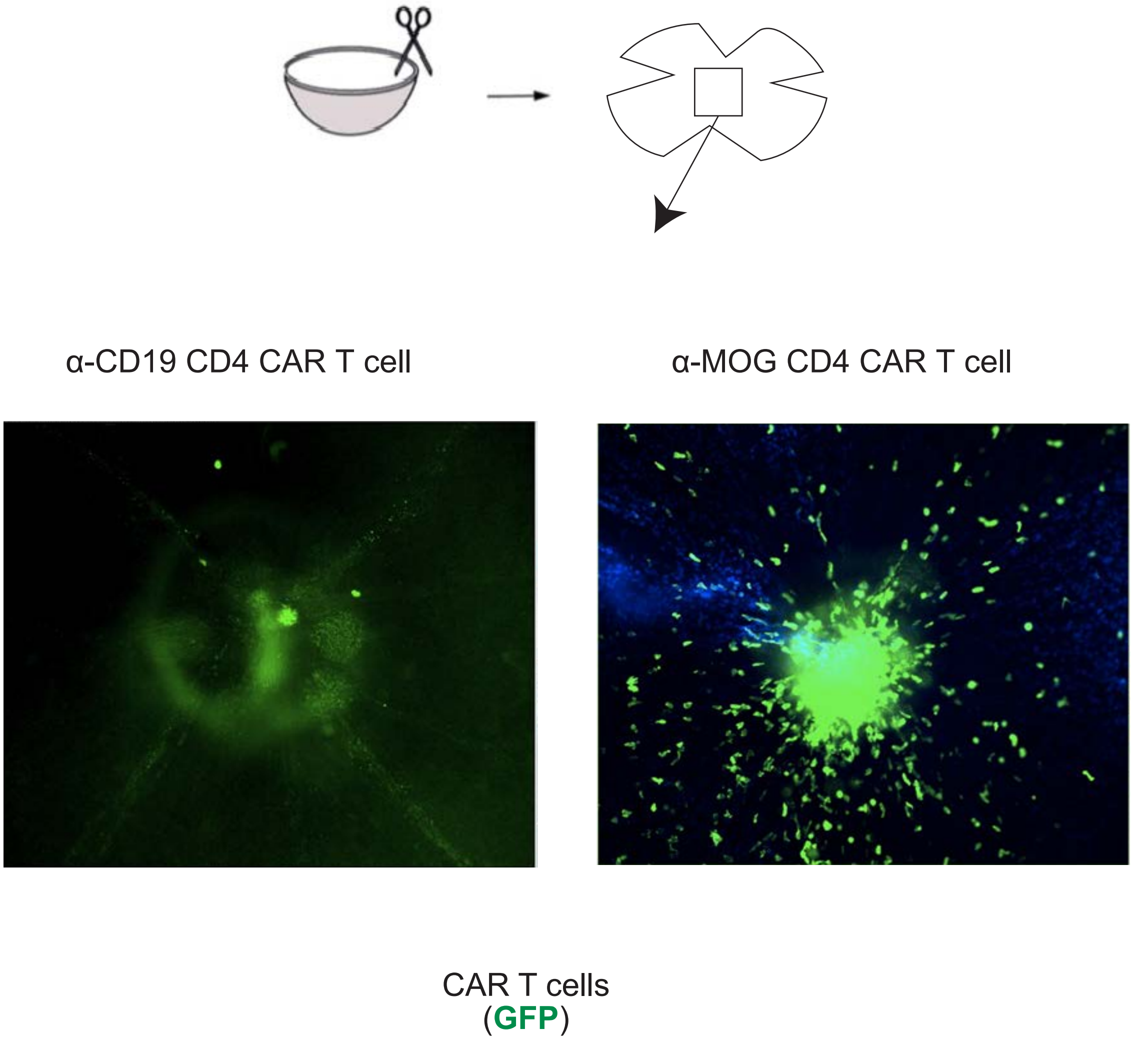


**Extended Data Fig. 5: Human α-MOG CAR CD4+ T cells infiltrate the retina (related to Fig. 4).**
Human α-MOG CAR CD4+ T cells infiltrate the mice's retina. NSG mice were injected with 5 x $10^6$ primary human CD4+ T cells expressing α-MOG CAR or control α-CD19 CAR. All the T cells were also engineered to express GFP constitutively. Around two weeks later, the retinas were collected and flat-mounted. GFP signal was measured to assess T cell localization. α-MOG CAR CD4+ T cells resulted in infiltration of the retina. Representative images.

**Extended Data Fig. 6: Human α-MOG CAR CD4+ T cells infiltrate the optic nerve (related to Fig. 5)**

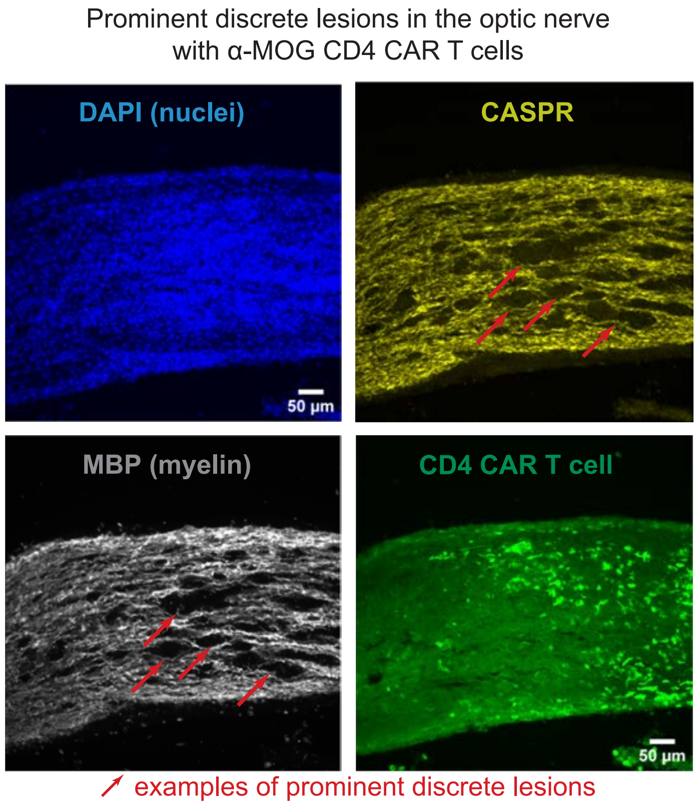


**Extended Data Fig. 6: Human α-MOG CAR CD4+ T cells infiltrate the optic nerve (related to Fig. 5).**
Human α-MOG CAR CD4+ T cells infiltrate the mice's optic. NSG mice were injected with 5 x $10^6$ primary human CD4+ T cells expressing α-MOG CAR and GFP constitutively. Two weeks later, the optic nerves were collected and stained for DAPI (nuclei), CASPR (paranodes), and MBP (myelin). α-MOG CAR CD4+ T cells infiltrated the optic nerve (GFP) and lead to prominent discrete demyelination (red arrow), like in human optic neuritis. Note that these images are the same as in Fig. 5A. Representative images.

**Extended Data Fig. 7: Clemastine protects the optic nerve myelin from human α-MOG CAR CD4+ T cells (related to Fig. 6)**

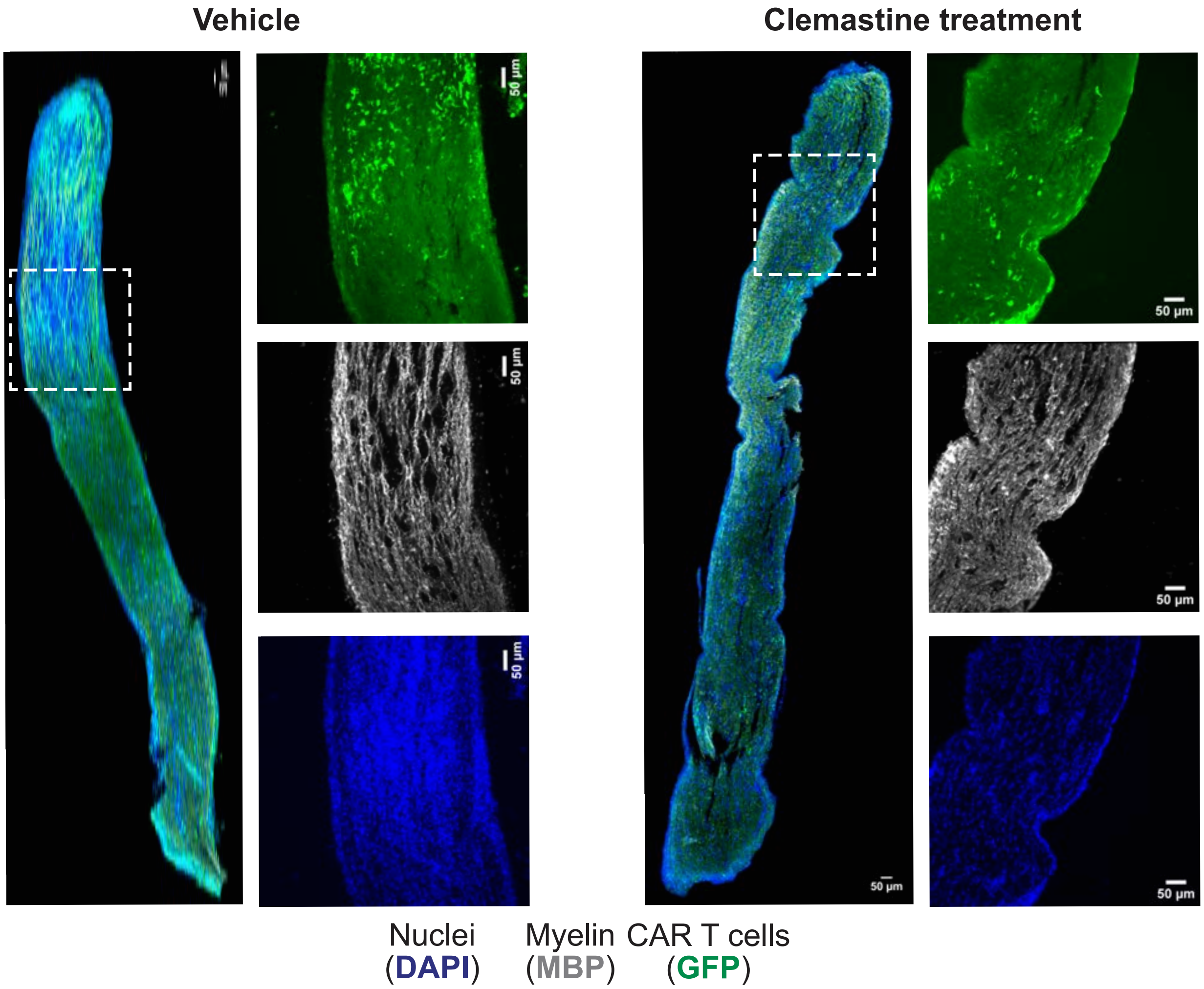


**Extended Data Fig. 7: Clemastine protects the optic nerve myelin from human α-MOG CAR CD4+ T cells (related to Fig. 6).**
Clemastine protects from the demyelination caused by the human α-MOG CAR CD4+ T cells. Primary human CD4+ T cells were engineered to express α-MOG CAR and GFP-2A-effLuc. 5 x $10^6$ primary human CD4+ T cells were injected i.v. in immunocompromised NSG mice and treated with vehicle (n=6) or clemastine (n=8) by daily gavage starting at day one post T cell injection. The optic nerves were processed by immunofluorescence and stained for MBP (myelin) and CASPR (paranodes). CAR T cells were engineered to express GFP and can be localized that way. Clemastine-treated optic nerves had more myelin (MBP stain) and more paranodes (CASPR stain) despite having α-MOG CAR T cells present.

**Extended Data Fig. 8: Clemastine does not affect human α-MOG CAR CD4+ T cells activation and ability to secrete pro-inflammatory cytokine when co-cultured with MOG⁺ K562s (related to Fig. 6)**

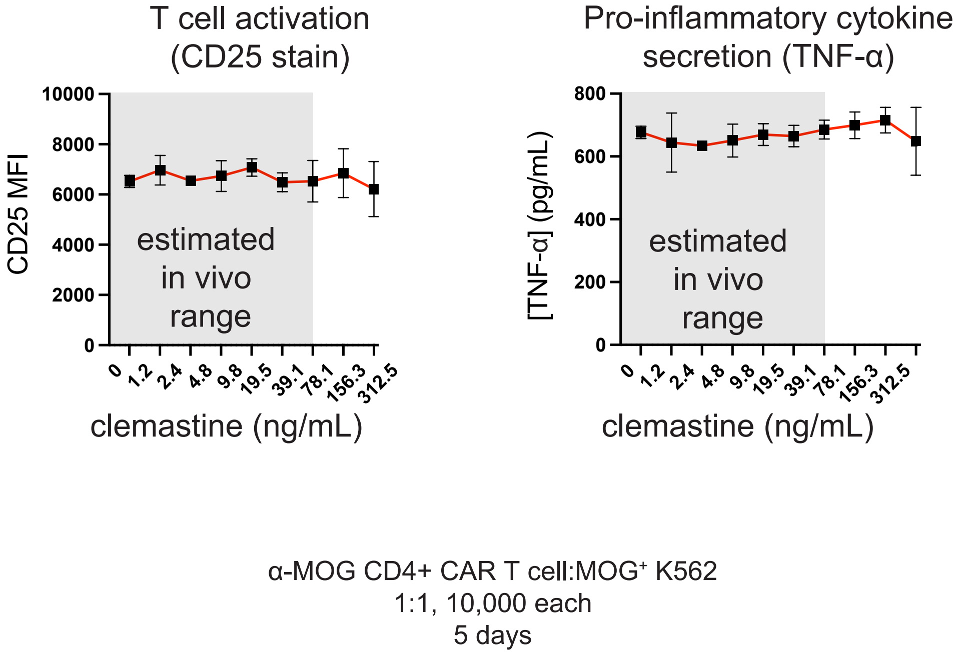


**Extended Data Fig. 8: Clemastine does not affect human α-MOG CAR CD4+ T cells activation and ability to secrete pro-inflammatory cytokine when co-cultured with MOG⁺ K562s (related to Fig. 6).**

Activation and pro-inflammatory cytokine secretion in vitro. Primary human CD4+ T cells transduced with α-MOG CAR were co-cultured with K562 expressing MOG at indicated clemastine concentrations, E:T ratio, duration, and cell numbers. Activation state was assessed by flow cytometry using a CD25 stain. Secretion of pro-inflammatory cytokines TNF-α was assessed by ELISA from the supernatant. n = 3, mean +/- std. The grey area shows the estimated in vivo concentration range for clemastine-treated mice.

**Extended Data Table 1: Human α-MOG CAR CD4+ T cells reproducibly induce severe optic neuritis.** Histology analysis of mice treated with α-MOG CAR CD4+ T cells by immunofluorescence. Summary of different experiments.

| Analysis | Optic neuritis |
|---|---|
| **Immunofluorescence** | 9/9 (100%) |
